# Towards the automated detection of interictal epileptiform discharges with magnetoencephalography

**DOI:** 10.1101/2023.07.14.548995

**Authors:** Raquel Fernández-Martín, Odile Feys, Elodie Juvené, Alec Aeby, Charline Urbain, Xavier De Tiège, Vincent Wens

## Abstract

The analysis of clinical magnetoencephalography (MEG) in patients with epilepsy traditionally relies on the visual identification of interictal epileptiform discharges (IEDs), which is time consuming and dependent on (subjective) human criteria. Data-driven approaches enabling both spatial and temporal localization of epileptic spikes would represent a major leap forward in clinical MEG practice. Here, we explore the ability of Independent Components Analysis (ICA) and Hidden Markov Modeling (HMM) to automatically detect and localize IEDs. Combined with kurtosis mapping, we developed a fully automated identification of epileptiform independent components (ICs) or HMM states. We tested our pipeline on MEG recordings at rest from 10 school-age children with either focal or multifocal epilepsy and compared results with the traditional MEG analysis performed by an experienced clinical magnetoencephalographer. In patients with focal epilepsy, both ICA- and HMM-based pipelines successfully detected visually identified IEDs with high sensitivity, but also revealed low-amplitude IEDs unidentified by the visual detection. Success was more mitigated in patients with multifocal epilepsy, as our automated pipeline missed IED activity associated with some foci—an issue that could be alleviated by *post-hoc* manual selection of epileptiform ICs or HMM states. Therefore, IED detection based on ICA or HMM represents an efficient way to identify spike localization and timing, with heightened sensitivity to IEDs compared to visual MEG signal inspection and requiring minimal input from clinical practitioners.

## 1. Introduction

Epilepsy is a disease of the brain characterized by at least two unprovoked seizures, a high probability of recurrence after a first seizure, or by a diagnosis of an epilepsy syndrome (Fisher et al., 2005, 2014). Epileptic seizures can be classified as focal when the seizure onset is localized or more widely distributed within one hemisphere, or as generalized when they rapidly engage bilateral networks. The onset of focal seizures may be characterized by seizure semiology and by non-invasive electrophysiological recordings such as scalp electroencephalography (EEG) or magnetoencephalography (MEG). Although MEG/EEG recordings of epileptiform discharges during clinically observed seizures (i.e., ictal epileptiform discharges) may be difficult (Alkawadri et al., 2018), they allow the investigation of subclinical events such as epileptiform discharges occurring between the seizures (i.e., interictal epileptiform discharges (IEDs)) (Kirsch et al., 2006; Kural et al., 2020; Ossadtchi et al., 2004; Tatum et al., 2018). In a majority of the patients, IEDs occur far more often than seizures, have a better signal to noise ratio than ictal epileptiform discharges (Alkawadri et al., 2018) and contribute to the non-invasive delineation of epileptogenic zone (i.e., the smallest area of the brain that can generate epileptic seizures) (Lüders et al., 2016; Nair & Beaumont, 2006). Thus, MEG/EEG recordings of IEDs represent an invaluable diagnostic tool in patients with epilepsy. In particular, MEG—which combines high temporal and good spatial resolutions—brings an established clinical added value to the non-invasive localization of the irritative zone (i.e., the brain area generating IEDs), and thus to the presurgical evaluation of patients with refractory focal epilepsy (Burgess, 2020; Chowdhury et al., 2015; De Tiège et al., 2012; Feys, et al., 2022b; Krishnan et al., 2015; Kural et al., 2020; Nissen et al., 2017; Rampp et al., 2019) especially in perisylvian epilepsy (Feys et al., 2023).

The predominant, clinically validated approach to identify IEDs in MEG signals combines the visual identification of IEDs within magnetic sensor signal traces with the source localization of their corresponding neural network by fitting a single or multiple equivalent current dipoles (ECDs) to spike events (Bagic et al., 2011; Ebersole, 1997). This process remains unavoidably time consuming (on average ∼8h (De Tiège et al., 2017)), demands both clinical and procedural expertise, and ultimately depends on subjective interpretation (notwithstanding its success and ongoing efforts to establish a common practice (Bagic et al. 2011; Laohathai et al., 2021)). Alternatively, kurtosis mapping based on beamformer spatial filtering has been utilized to automatically localize sources of IED activity without explicit identification of spike events (Hall et al., 2018; Kirsch et al., 2006; Robinson et al., 2004). However, kurtosis mapping may fail in situations of highly frequent epileptic spikes, leading to lower kurtosis values and undetected IED sources (Seedat et al., 2022). Further, kurtosis does not provide temporal information necessary to examine single IEDs or to characterize spike propagation dynamics. The development of automatic, data-driven methods to characterize both the spatial localization and the temporal dynamics of IEDs has been undertaken using different methodologies. Independent Component Analysis (ICA) represents a natural candidate given its ability to isolate sparse, high-amplitude events such as the cardiac artefact (Vigário, 1999) and epileptic spikes as demonstrated in EEG (Hoeve et al., 2003; James et al., 1997) and in MEG (Malinowska et al., 2014; Pizzo et al., 2019). In particular, Ossadtchi et al. identified independent components (ICs) of MEG data that exhibit spike-like characteristics (Ossadtchi et al., 2004). They subsequently supplemented this analysis with Hidden Markov Modeling (HMM) of spike events to estimate patterns of spike propagation (Ossadtchi et al., 2005), and spatio-temporal event clustering via convolutional sparse coding to identify IEDs and locate the seizure onset zone (Chirkov et al., 2022). Multivariate implementations of HMM have also been proposed for data-driven IED detection in MEG recordings (Seedat et al., 2022; Ye et al., 2022). These implementations were originally developed to model functional brain network activations as discrete brain states and come in two distinct versions: one focusing on MEG amplitudes (amplitude HMM, aHMM; Baker et al., 2014) and another relying on time-embedded MEG signals (time-embedded HMM, TE-HMM; Vidaurre et al., 2018). Seedat et al. (2022) used TE-HMM to infer epileptic network states from MEG data, though their identification amongst physiological network states required visual selection by a clinician (Seedat et al., 2022). This TE-HMM approach was further combined with a graph-theoretical analysis of functional connectivity to identify pathological states (Ye et al., 2022).

To the best of our knowledge, the ability of aHMM to disclose epileptic networks has not been tested yet. A comparison of the ICA and HMM frameworks is also lacking, as well as their benchmarking against kurtosis mapping (Hall et al., 2018) and visual IED detection following traditional clinical practice (Laohathai et al., 2021).

Here, we sought to design, test, and compare two distinct pipelines for IED detection and spatio-temporal localization from resting-state MEG data of individual patients with (multi)focal epilepsy, with minimal-to-no input from clinical magnetoencephalographer. First, we consider IED detection based on ICA (Ossadtchi et al., 2004) supplemented with a univariate, two-state aHMM of IC time courses for automated clustering of IED events (in the spirit of (Chirkov et al., 2022) but relying on aHMM rather than convolutional sparse coding). Second, we examine the ability of multivariate aHMM to detect IED activity (somewhat similarly to (Seedat et al., 2022; Ye et al., 2022), but focusing on MEG amplitudes rather than time-embedded signals). The main rationale to choose aHMM being that IEDs are primarily characterized by pathologically high-amplitude events (compared to physiological events such as oscillatory power bursts; see, e.g., (Coquelet et al., 2022)).

Within both frameworks, epileptiform ICs or aHMM states are identified automatically by comparing their spatial distribution with kurtosis mapping. Further, we compare the outcomes with visual IED detection performed by an experienced clinical magnetoencephalographer blind to the data-driven classifications. We hypothesized that both ICA and HMM-based pipelines would allow to recover visually detected IEDs in a fully automated manner.

## 2. Methods

### 2.1 Participants

Table 1 details patients’ clinical characteristics and diagnosis.

**Table 1.** Abbreviations. F: female; M: male; EE-CSWS: epileptic encephalopathy with continuous spike-and-wave during sleep; SeL-ECTS: self-limited epilepsy with centro-temporal spikes; RFE: refractory focal epilepsy; R: right; L: left; CT: centro-temporal; OT: occipito-temporal; O: occipital.

| **Patient** | **Age (y)** | **Sex** | **Type of Epilepsy** | **Scalp Location** |
| --- | --- | --- | --- | --- |
| 1 | 10 | F | Structural epilepsy | R-CT |
| 2 | 11 | M | EE-CSWS | R-OT |
| 3 | 8 | M | SeL-ECTS | CT-bilateral |
| 4 | 9 | F | SeL-ECTS | L-CT |
| 5 | 10 | M | SeL-ECTS | L,R-CT and R-O |
| 6 | 10 | M | SeL-ECTS | R-CT |
| 7 | 11 | M | SeL-ECTS | L-CT |
| 8 | 5 | F | SeL-ECTS | L-CT |
| 9 | 9 | F | SeL-ECTS | L-CT |
| 10 | 11 | F | Structural RFE | R-CT |

We included 10 children with focal epilepsy (5 females and 5 males; age range: 5-11 years) and frequent IEDs on previous scalp EEG. The majority of them suffered from self-limited epilepsy with centro-temporal spikes (SeLECTS) (8 patients; 1 with epileptic encephalopathy with continuous spike-and-wave during sleep (EE-CSWS)). Patient 1 has periventricular nodular heterotopias adjacent to the frontal horn of the left lateral ventricle and a nonspecific hyperintensity of the right frontal white matter on T2-wheighted MRI. Patient 10 has a right anterior temporal post-resection cavity following surgical ablation of a dysembryoplastic neuroepithelial tumor.

All participants and their legal representative signed a written informed consent approved by the Hôpital Erasme Ethics Committee.

### 2.2 Data acquisition

All patients underwent at least 5 minutes of resting-state MEG activity. Data were recorded at the Hôpital Erasme using a 306-channel whole-scalp neuromagnetometer installed in a lightweight magnetically shielded room (Neuromag Triux and Maxshield; MEGIN Oy, Helsinki, Finland; see (De Tiège et al., 2008) for details) with analog band-pass at 0.1-330 Hz and sampling frequency at 1 kHz. Patients sat comfortably in the MEG armchair with their eyes open and did not suffer from clinical or electrical seizures during the recording. Four coils were used to track and eventually correct for head movements. Their position relative to fiducials (nasion and tragi) was digitized beforehand using an electromagnetic tracker (Fastrack, Polhemus, Colchester, Vermont, USA), along with at least 200 face and scalp points for subsequent co-registration with structural brain magnetic resonance imaging (MRI). The latter was acquired in each patient using a high-resolution 3D T1-weighted MRI (Intera, Philips, The Netherlands) after the MEG recording.

### 2.3 Data preprocessing and source projection

Each MEG data was denoised using signal space separation (Taulu et al., 2005) with head movement correction (Maxfilter v2.2.14, MEGIN Oy; with default parameters and no bad channel detected in the process). We filtered the data in the 0.5-45 Hz frequency band and applied a first ICA (FastICA algorithm with dimension reduction to 30 and contrast nonlinearity *tanh*) to identify cardiac and eye movements artifacts (Vigário, 1999). Artefactual ICs were regressed out of the full-rank data (number of ICs removed per patient: 1 - 4, range). The resulting sensor data were filtered between 4 and 30 Hz.

Forward models were computed using the one-layer Boundary Element Method implemented in the MNE-C suite (Gramfort et al., 2014) based on each patient’s MRI segmented using the FreeSurfer software (Fischl, 2012). The source space was built by placing three orthogonal unit dipoles on a 5-mm cubic lattice sampling the MRI brain volume. The lattice was first defined in the Montreal Neurological Institute (MNI) template MRI volume and then non-linearly deformed onto each patient’s MRI with the Statistical Parametric Mapping software (SPM12) (William Penny, 2006). Neural source activity was reconstructed using Minimum Norm Estimation (MNE; Dale et al., 1993) with noise standardization for depth bias correction (Pascual-Marqui, 2002) based on the corresponding forward model, a noise covariance matrix estimated from empty-room MEG recordings (preprocessed with signal space separation and filtered in 4-30 Hz band), and the regularization parameter set according to a consistency condition (Wens et al., 2015). Each three-dimensional source signal was further projected on the direction of maximal power.

These cleaned MEG data and their source projection were then inputted to the two IED detection pipelines represented schematically in Fig. 1 and described hereunder.

**Figure 1.**
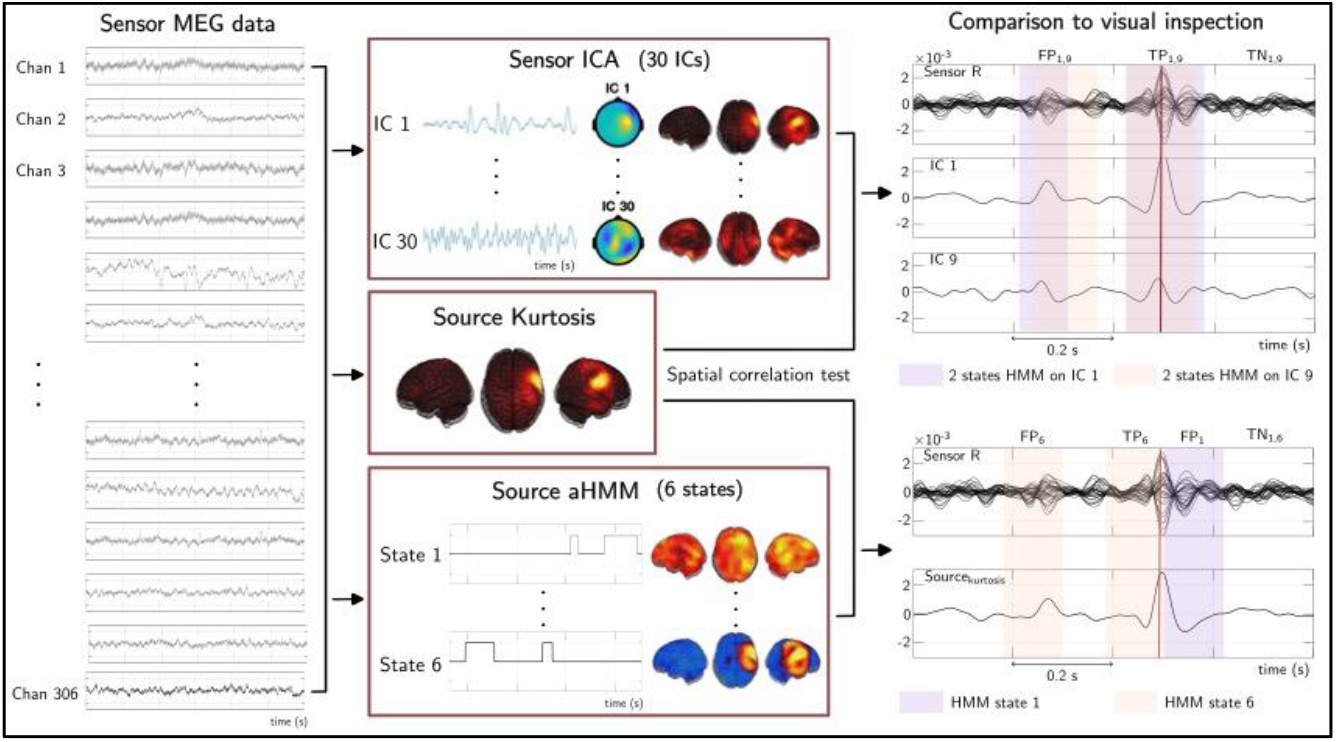
Schematic representation of the two pipelines proposed to localize IED spatially and temporally from sensor MEG data (**left**). These data are fed either to an ICA yielding 30 ICs characterized by a continuous time course, a sensor topography, and its source projection (**middle, top panel**) or to an aHMM of source amplitudes yielding 6 states characterized by state activation windows and source power maps (**middle, bottom panel**). In both cases, source maps are correlated spatially with source kurtosis mapping (**middle, central panel**) in order to select “epileptiform” ICs or aHMM states reflecting IED activity. Signal periods containing IEDs are then detected automatically (**right**, shaded windows), using either a two-state aHMM of the epileptiform ICs (**right, top panel**) or the epileptiform aHMM state activation windows (**right, bottom panel**). Comparison to visual inspection by a clinician was performed by labelling each data period according to whether or not visually detected IED (vertical red lines) occur in or out of activation windows (**right**; TP, true positive; FP, false positive; TN: true negative; FN: false negative; see main text for definitions). We show on the right the sensor data used for visual inspection along with the source with highest kurtosis. In this example, two epileptiform ICs were detected (IC1 and IC9; **right, top panel**) and two epileptiform aHMM states (states 1 and 6; **right, bottom panel**). Note that activation windows of two ICs may overlap but those of two aHMM states may not. In each case, two disjoint activations appear: the first does not coincide with visually detected spike (FP) but the second does (TP).

### 2.4 Independent Components Analysis and amplitude Hidden Markov Modeling

In our first pipeline, the exact same ICA than described in section 2.3. for artefact removal was applied again on the sensor MEG signals, this time in order to seek for IEDs (Ossadtchi et al., 2004). This allowed to decompose MEG data into 30 ICs, each characterized by a continuous activity time course and by a sensor topography (see “Sensor ICA” panel in Fig. 1). Spatial localization of brain sources involved in each IC was obtained by source projection of their scalp topography.

In our second pipeline, we applied aHMM to the projected source signals, closely following the approach described by Baker et al. (Baker et al., 2014) with two main differences. First, source projection was performed with MNE ((Coquelet et al., 2022); for applications, see also (Coquelet et al., 2020; Naeije et al., 2021; Puttaert et al., 2020; Roshchupkina et al., 2022)). Second, HMM state inference was performed at the single-subject level rather than the group level. Briefly, the amplitude time courses of source signals were estimated by Hilbert transformation, downsampled at 40 Hz with low-pass filtering at 10 Hz via moving-window averaging, demeaned, and normalized by the individual standard deviation, and finally dimensionally reduced to their 40 first principal components. The resulting data was fed into a 6-state HMM with Gaussian multivariate observation model along with the Viterbi algorithm (Rezek et al., 2005) as implemented in the HMM-MAR toolbox (Vidaurre et al., 2016). Each state was eventually characterized by a binary time series of the most probable state activation under the constraint that all states are temporally exclusive (see “Source aHMM” panel in Fig. 1). The spatial distribution of each state was characterized by a brain map showing regional source power increases/decreases upon state activation/deactivation. It was computed as the partial correlation between binary state activation time series and source amplitude signals.

### 2.4 Automated spike detection

To identify ICs and aHMM states related to IED activity, we considered the spatial correlation between brain maps associated with each IC/aHMM state and kurtosis mapping (see “Source Kurtosis” panel in Fig. 1). Each individual kurtosis map was obtained by imaging the kurtosis statistic of each projected source signal (Kirsch et al., 2006; Robinson et al., 2004), so regions with large values compared with physiological brain activity are indicative of temporally sparse, high-amplitude events presumed to reflect IEDs (Hall et al., 2018). Statistical significance of each spatial correlation was established according to two-tailed parametric correlation test at *p*<0.05 under the null hypothesis that Fisher-transformed spatial correlations follow a Gaussian distribution with mean zero and standard deviation (SD) 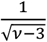 where *v* the number of spatial degrees of freedom estimated from the rank of the forward model (Coquelet et al., 2022). Significantly positive correlations allowed to identify “epileptiform ICs/aHMM states” as they co-localize with high-kurtosis regions. Of note, in most of the cases this may lead to more than one epileptiform IC and more than one epileptiform aHMM state.

We then relied on the time course of epileptiform ICs/aHMM states to detect the timing of IEDs. The binary character of aHMM state time courses directly enabled such temporal localization automatically (see right bottom panel in Fig. 1), but extra steps were required with ICA due to the continuous character of IC time courses (Chirkov et al., 2022). Here, we applied a 2-state HMM on the amplitude of each epileptiform IC (using a pipeline similar to the one described in section 2.3) in order to cluster time points into a high-amplitude state presumed to contain IEDs (selected as the HMM state with largest mean amplitude) and a low-amplitude state devoid from IEDs (i.e., the complementary HMM state). In this way, each epileptiform IC was associated to a binary IED detection time course similarly to the aHMM pipeline (see right top panel in Fig. 1). In cases where spatial correlation with kurtosis mapping revealed multiple epileptiform ICs or multiple epileptiform aHMM states, we gathered the corresponding binary IED detection time courses into a single “global” IED detection time course built as their logical union (i.e., detecting periods containing at least one IED detection event). This was used for IED counting purposes and quantitative comparison to visual IED detection.

### 2.5 Comparison to the visual IED identification

We further compared the IED detection efficiency of these two methods with visual detection. For this purpose, one clinical magnetoencephalographer (O.F.) analyzed sensor-level MEG signals, blindly to the automated analyses, according to well-established practice (Bagic et al., 2011; Laohathai et al., 2021) and marked the timing of each visually detected sensor-IED (VDS-IED). These VDS-IEDs were localized using ECD (source modelling software Xfit, MEGIN). Then, we defined several metrics allowing to compare VDS-IEDs and IED detection periods revealed by the global detection events of epileptiform ICs or aHMM states: IED detection rate (IED-DR), sensitivity and specificity. Of note, these three metrics use VDS-IEDs as the “ground truth”, i.e., they rely on the hypothesis that VDS-IEDs identify all IEDs correctly, a hypothesis that we then re-assessed using our data (see below).

We measured the IED-DR as the percentage of the number (#VDS-IED_ICA/aHMM_) of VDS-IEDs that occurred within the IED detection periods outputted by our ICA or aHMM pipelines:

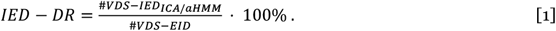

We also estimated the sensitivity and the specificity of our ICA and aHMM pipelines, according to standard expressions:

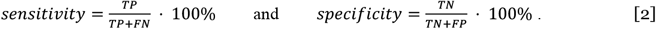

Here, TP stands for the number of IED detection periods that contain at least one VDS-IED (i.e., interpreted as true positives assuming that VDS-IEDs correctly identify all IEDs), FP the number of IED detection periods that do not contain any VDS-IED (false positives), FN the number of periods without IEDs according to our global detection time course that contain at least one VDS-IED (false negative), and TN the number of periods without IEDs according to our global detection time course that contain no VDS-IED (true negative). See Fig.1 (right) for examples. In this framework, high sensitivity for our data-driven pipelines would mean that they automatically recovered a large fraction of IEDs detected by the clinical magnetoencephalographer, whereas low sensitivity would reveal that they missed many VDS-IEDs. High specificity would mean that most automated IED detection events do correspond to VDS-IEDs, whereas low specificity would mean that many events were not marked as VDS-IED by the clinician. Of notice, IED-DR and sensitivity are closely related but different: the former takes into account the possibility that several VDS-IEDs occur within a single IED detection event, whereas the latter neglects this aspect.

To correctly interpret the above metrics, we further examined the validity of our assumption that VDS-IEDs correctly identify all IEDs. To that aim, we reconstructed brain power maps associated to each IED detection period (i.e., source-projected signal power map, time averaged within the corresponding period), and we averaged all TP events and all FP events separately to generate a TP map and a FP map. Under our assumption about VDS-IEDs, the FP map would not correspond to IEDs and thus should have both lower power values and a spatial distribution very different from the TP map (that should locate IED activity).

## 3. Results

The maximum source kurtosis value in each of the 10 patients was well above typical values in healthy subjects (i.e., slightly above the Gaussian value *kurt*=3 (Karl Pearson, 1905); Table 2), attesting that all of them exhibited IEDs in their sensor-level MEG signals. This was confirmed independently by visual inspection of the clinical magnetoencephalographer.

**Table 2.** Maximum values of source kurtosis at the source level across patients.

| *Patient* | **1** | **2** | **3** | **4** | **5** | **6** | **7** | **8** | **9** | **10** |
| --- | --- | --- | --- | --- | --- | --- | --- | --- | --- | --- |
| *Max(kurt)* | 15.6 | 11.1 | 42.1 | 35.7 | 27.6 | 27.8 | 56.8 | 34.6 | 11.4 | 11.7 |

We describe the outcomes of our ICA- and aHMM-based methods in two separate sections. In each case, we report summary statistics (IED-DR, sensitivity and specificity; see Table 3 and Table 4) and discuss two representative patients—one showing good performance of the automated IED detection pipeline, and another showing mitigated performance. In the latter case, we further sought to explain where the automated IED detection failed and to provide a solution.

**Table 3.** IED detection rate (IED-DR), sensitivity and specificity of the ICA-based pipeline for each patient.

| **Patient** | **1** | **2** | **3** | **4** | **5** | **6** | **7** | **8** | **9** | **10** |
| --- | --- | --- | --- | --- | --- | --- | --- | --- | --- | --- |
| **IED-DR** | 99.2% | 75.0% | 76.3% | 96.3% | 55.9% | 96.4% | 100% | 99.0% | 100% | 98.8% |
| **Sensitivity** | 99.2% | 77.8% | 75.7% | 96.3% | 63.2% | 96.3% | 100% | 98.9% | 100% | 98.7% |
| **Specificity** | 54.8% | 50.2% | 52.3% | 58.1% | 52.2% | 52.5% | 50.4% | 53.0% | 50.9% | 53.3% |

**Table 4.** Spike detection rate (IED-DR), sensitivity and specificity of the aHMM-based pipeline for each patient.

| **Patient** | **1** | **2** | **3** | **4** | **5** | **6** | **7** | **8** | **9** | **10** |
| --- | --- | --- | --- | --- | --- | --- | --- | --- | --- | --- |
| *IED-DR* | 94.6% | 41.7% | 47.4% | 92.6% | 97.0% | 89.3% | 100% | 89.7% | 92.9% | 91.3% |
| *Sensitivity* | 93.9% | 42.9% | 54.6% | 92.6% | 96.3% | 95.7% | 100% | 88.4% | 92.9% | 91.4% |
| *Specificity* | 60.7% | 49.4% | 51.9% | 63.1% | 54.1% | 56.2% | 53.0% | 57.3% | 51.7% | 54.2% |

### 3.1 Automated spike detection based on Independent Component Analysis

Seven out of 10 patients exhibited both IED-DR and sensitivity above 96%, showing that ICA-based IED detection was able to identify a large majority of VDS-IEDs in an automated way. The 3 other patients had lower IED-DR/sensitivity (patients 2, 3, and 5; range 56%–78%) reflecting that a substantial fraction of VDS-IEDs were missed in these patients (patient 2, 25%; patient 3, 24%; patient 5, 54%; fraction computed as 100% − *SDR*). In all patients, specificity ranged between 50% and 58%, meaning that almost half IED events extracted from ICA did not correspond to VDS-IEDs and were labelled as FPs.

Figure 2 details the outcome of the ICA pipeline in one patient with high IED-DR/sensitivity (patient 1). Kurtosis mapping was compatible with a right centro-temporal irritative zone (Fig. 2(a)) and concurred with ECD fitting of VDS-IEDs (obtained independently by the clinical magnetoencephalographer; see Fig. 2(b). Figure 2(c) identifies two epileptiform ICs that co-localized with kurtosis mapping. We show 2 seconds of example data in Fig. 2(e) along with IED detection windows of these two epileptiform ICs (shaded areas) and VDS-IEDs (vertical red lines). This illustrates the high IED-DR/sensitivity (99.2%) since all VDS-IEDs occurred here inside IED detection windows, and its fairly low specificity (54.8%) since several IED detection windows occurred without any VDS-IED. Further examination of Fig. 2(e) shows that those windows devoid of VDS-IED (termed FP in our terminology) contain spike-like events of lower amplitude than those successfully identified by the clinical magnetoencephalographer (TP). This suggests that our ICA pipeline allows to identify small-amplitude IEDs that were not recognized during visual inspection. To confirm this hypothesis, we mapped source activations in TP events and in FP events (Fig. 2(d)). Unsurprisingly, the TP maps in both epileptiform ICs co-localized with kurtosis mapping and exhibited high (noise-normalized) peak amplitude. Source activation in the FP maps also co-localized with kurtosis mapping albeit with peak amplitude 40%–50% lower. This provides evidence that FP events could correspond to genuine IEDs that were not successfully identified by visual detection due to their low amplitude, which was similar to background noise. These observations generalized to all patients.

**Figure 2.**
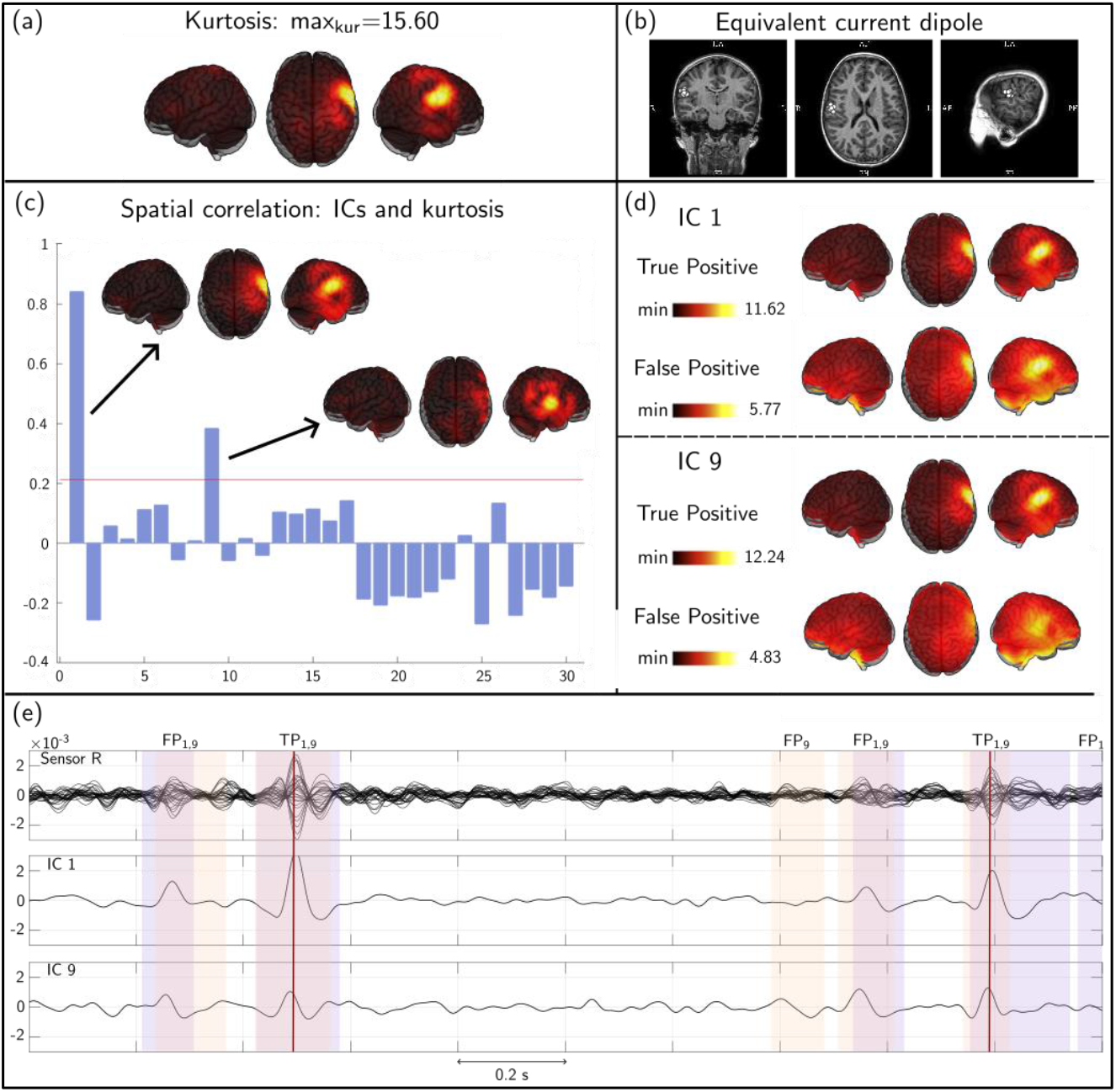
ICA-based IED detection in patient 1. (a) Source kurtosis mapping. The peak value is indicated (see also Table 2). (b) Equivalent current dipole (ECD) analysis of VDS-IEDs obtained independently by an experienced clinician. The MRI is shown here in radiological convention. (c) Spatial correlation between each IC source map and the kurtosis map. Epileptiform ICs correspond to ICs with significantly positive correction (statistical threshold represented with a horizontal red line). In this case, IC 1 and IC 9 were identified as epileptiform. (d) Source activation maps associated with “True Positive” detection events (TP, containing VDS-IEDs) and “False Positive” events (FP, devoid of VDS-IEDs). Color scales report noise-normalized source amplitude. Note the difference in scale maxima between TP and FP cases. (e) Two seconds of example times courses. Upper part, MEG data of right-hemisphere sensors; middle and lower parts, epileptiform ICs. The shaded windows show the IED detection events built using a 2-state aHMM of each epileptiform IC time course (IC 1, violet; IC 9, orange). Vertical lines indicate VDS-IEDs.

Figure 3 depicts the outcome of the ICA pipeline in one of the 3 patients showing substantially lower IED-DR/sensitivity (patient 5; IED-DR and sensitivity ≤ 63%). Kurtosis mapping (Fig. 3(a)) and ECD analysis of VDS-IEDs (Fig. 3(b)) revealed epileptiform foci in both right-occipital and right centro-temporal regions. The ECD analysis also highlighted a few left centro-temporal VDS-IEDs. Only one epileptiform IC localizing the right occipital focus was identified automatically by spatial correlation with kurtosis (Fig. 3(c)). This IC missed several IEDs that were clearly identified by visual inspection. Given the presence of other foci in this patient, we took a step back, inspected all ICs, and identified another IC exhibiting clear epileptiform activity at the right centro-temporal focus (Fig. 4). Including both the right-occipital IC (obtained by spatial correlation) and the right centro-temporal IC (obtained by visual inspection) allowed to increase the IED-DR from 56% to 99% and sensitivity from 63% to 98%, so these two ICs accounted for the large majority of VDS-IEDs, while keeping specificity to 54%. Therefore, supplementing our ICA pipeline with a visual inspection of ICs instead of the fully automated kurtosis correlation test, allowed to recover results similar to the seven other patients.

**Figure 3.**
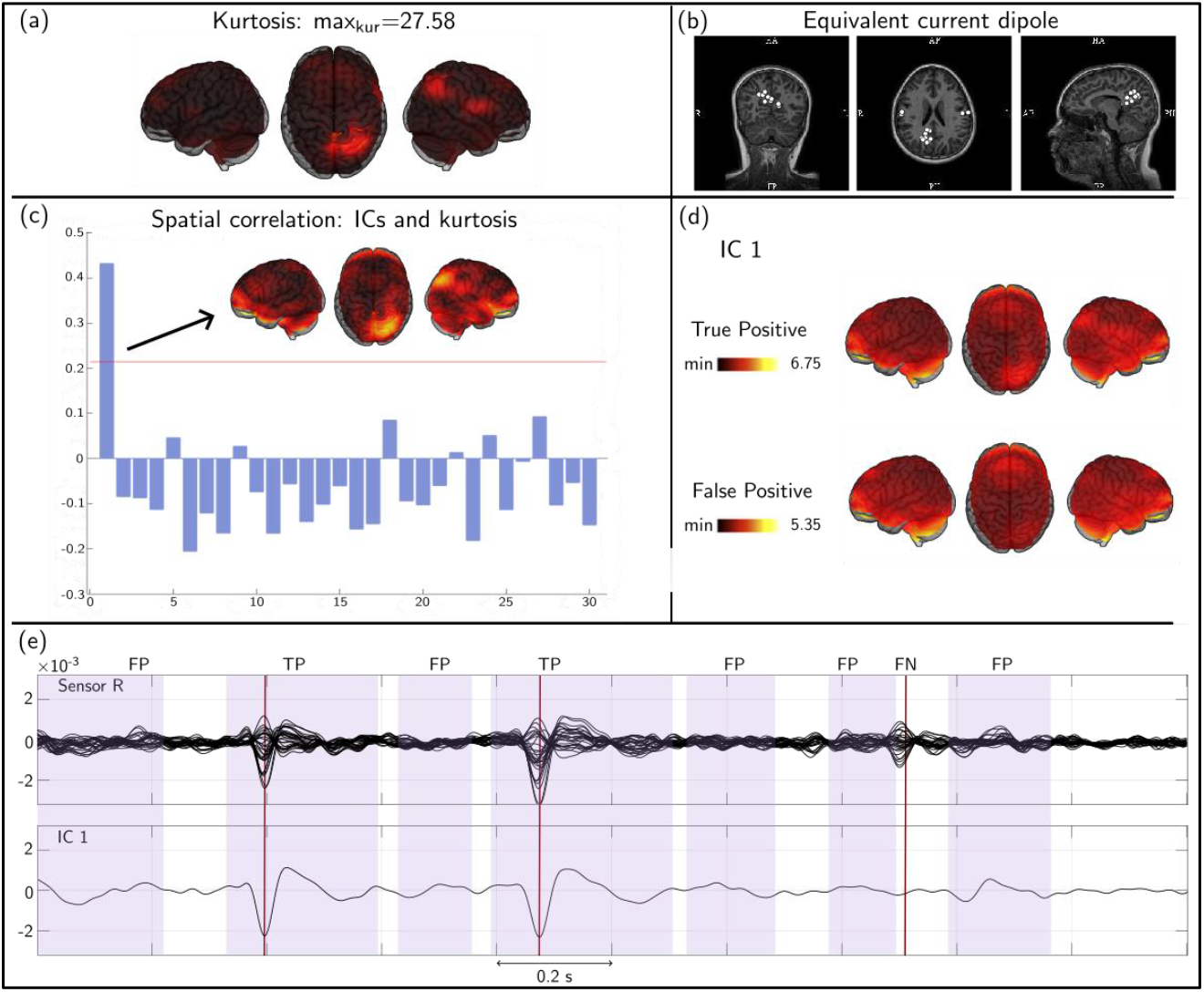
ICA-based IED detection in patient 5. All is as in Fig. 2. Here, IC 1 was the only component passing the kurtosis correlation test (panel (c)). The example in panel (e) shows one VDS-IED that was missed by IED detection events of IC 1 (shaded windows).

**Figure 4.**
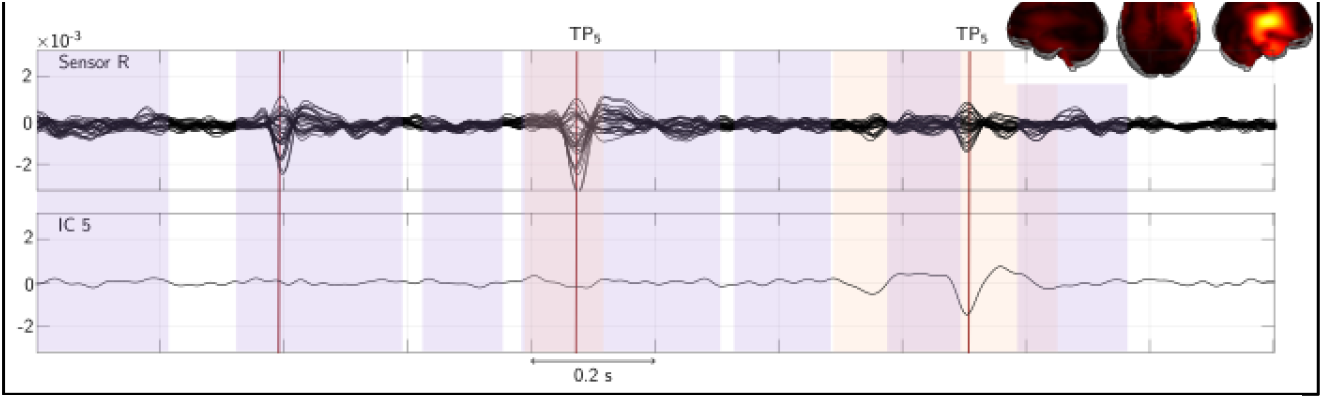
Visually selected IC 5 of patient 5. The time course of IC 5 is illustrated in the same 2-second period than in Fig. 3(e) along with IED detection events of IC 1 (purple shaded windows, see also Fig. 3(e)) and of IC 5 (orange shades). Vertical lines indicate VDS-IEDs. The source map of IC 5 is shown in the upper right.

The situation for patients 2 and 3 was similar. In fact, reviewing Table 3 in light of the above and of Table 1, it turns out that all 3 patients with initially low IED-DR/sensitivity suffered from multifocal epilepsy, whereas the 7 patients with high IED-DR/sensitivity had a single epileptic focus. In general, epileptiform ICs turned out to peak over a single epileptic focus, so that in the multifocal cases the spatial correlation with kurtosis (which exhibits multiple peaks) would miss some epileptiform ICs due to low correlation values. Fortunately, inspecting all IC maps allowed to find quickly missed epileptiform ICs and reach high IED-DR/sensitivity comparable to unifocal cases.

### 3.2 Automated spike detection based on Hidden Markov Modeling

The results for the aHMM-based pipeline followed a pattern very similar to those of the ICA-based pipeline. The IED-DR/sensitivity was above 88% for 8 out 10 patients and between 40% and 50% for the 2 other patients (patients 2 and 3), while specificity consistently ranged between 52% and 63% (Table 4). Of note, these statistics were not significantly different from those obtained with the ICA pipeline (Wilcoxon rank tests, *p*>0.16). The low specificity reflected the automated detection by epileptiform aHMM states of low-amplitude IED events left undetected by visual inspection, as was the case with ICA. Interestingly, patient 5 automatically resulted in high IED-DR/sensitivity with aHMM even though it was not the case with ICA due to the presence of multiple epileptic foci.

Figure 5 details the output of the aHMM-based IED detection in a patient with high IED-DR/sensitivity (patient 4). This patient exhibited a single epileptic focus in the left centro-temporal region (Fig. 5(b)) for kurtosis mapping; Fig. 5(c) for ECD fitting of VDS-IEDs). Among the 6 states inferred from aHMM of source-projected MEG data (Fig. 5(a)), only one was disclosed as “*epileptiform*” based on the spatial correlation test with kurtosis mapping. This state was sufficient to identify the large majority of VDS-IEDs (IED-DR/sensitivity, 93%; see Table 4 and Fig. 5(f) but could also identify lower amplitude IEDs (termed FP events) that were missed by visual inspection (see Fig. 5(f) and the similarity of TP and FP maps in Fig. 5(e), which explains the 63% specificity (Table 4).

**Figure 5.**
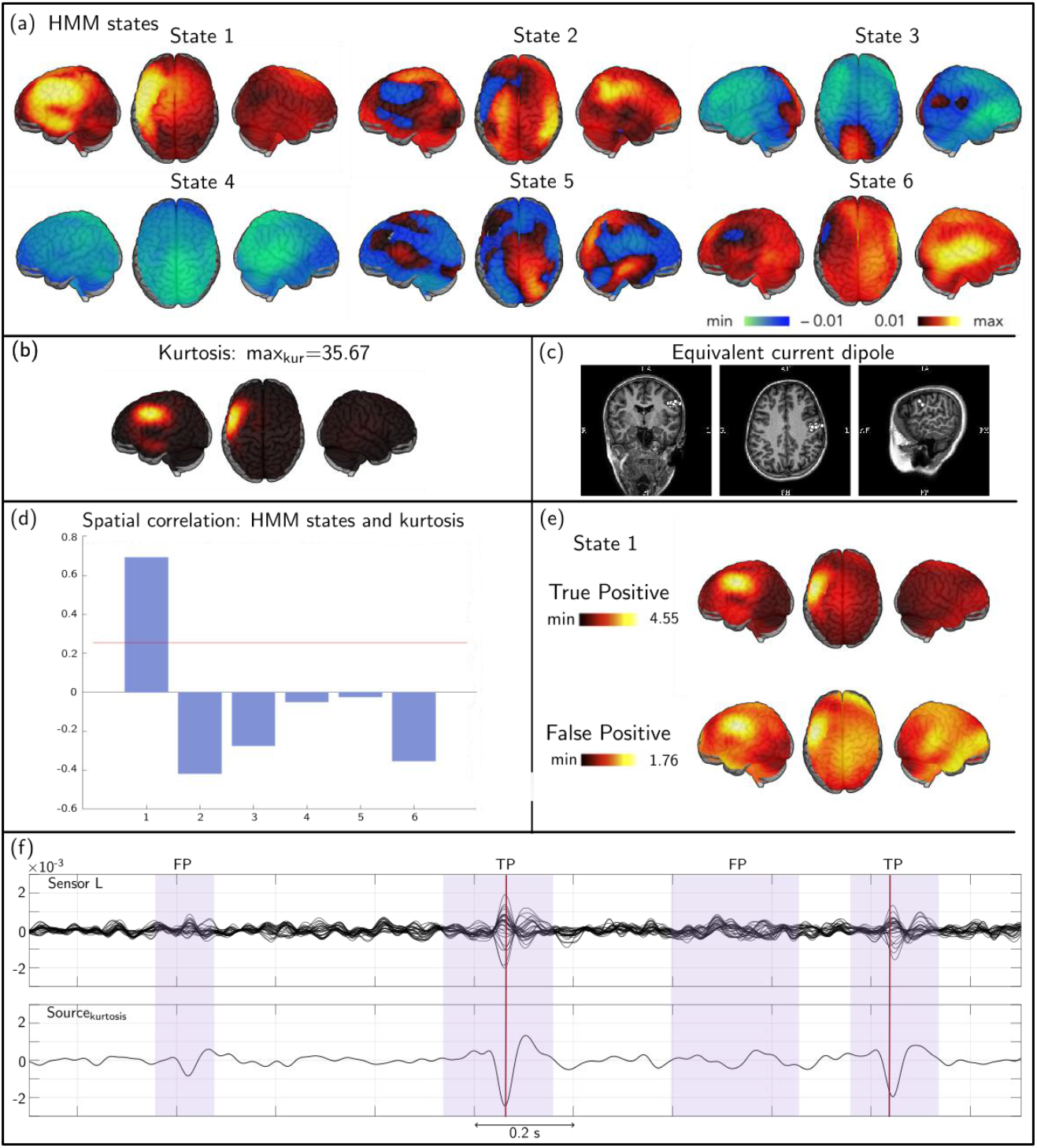
aHMM-based IED detection in patient 4. (a) State power maps of the 6-state aHMM. (b) Source kurtosis mapping. The peak value is indicated (see also Table 2). (c) Equivalent current dipole (ECD) analysis of VDS-IEDs obtained independently by an experienced clinician. The MRI is shown in radiological convention. (d) Spatial correlation between each state map and the kurtosis map. Epileptiform states correspond to aHMM states with significantly positive correction (statistical threshold represented with a horizontal red line). In this case, state 1 was identified as epileptiform. (e) Source activation maps associated with “True Positive” detection events (TP, containing VDS-IEDs) and “False Positive” events (FP, devoid of VDS-IEDs). Color scales report noise-normalized source amplitude. Note the difference in scale maxima between TP and FP cases. (f) Two seconds of example times courses. Upper part, MEG data of left-hemisphere sensors; lower part, maximum-kurtosis source signal. The shaded windows show the IED detection events (here, state 1 activation periods; violet shade). Vertical lines indicate VDS-IEDs.

A case with low IED-DR/sensitivity is illustrated in Fig. 6 (patient 3). This patient had a bilateral centro-temporal epilepsy (Fig. 6(b–c)). The automated kurtosis correlation test revealed two epileptiform states, both of which exhibiting power increases at the left focus, but missed IED activity emerging from the right focus, explaining the low IED-DR (47%) and sensitivity (55%). A glance at all 6 aHMM state maps allowed to identify one state with power increase above the right focus (state 1 in Fig. 6(a)) and checking its time course indeed confirmed that it detects IED events (Fig. 7). Its correlation with the kurtosis map was low because of the imbalance in kurtosis values at the two foci (Fig. 6(b)) and the anti-symmetry of its power map (Fig. 6(a)). Adding this visually selected state to the two automatically detected epileptiform states allowed to drastically increase IED-DR/sensitivity to 98% and slightly increased specificity to 64%. A similar improvement of aHMM-based detection statistics held in patient 2. As for the ICA pipeline, we conclude that the kurtosis correlation test may fail to identify automatically epileptiform states in patients with multifocal epilepsy, but this issue can be circumvented by a visual comparison of kurtosis and aHMM state maps.

**Figure 6.**
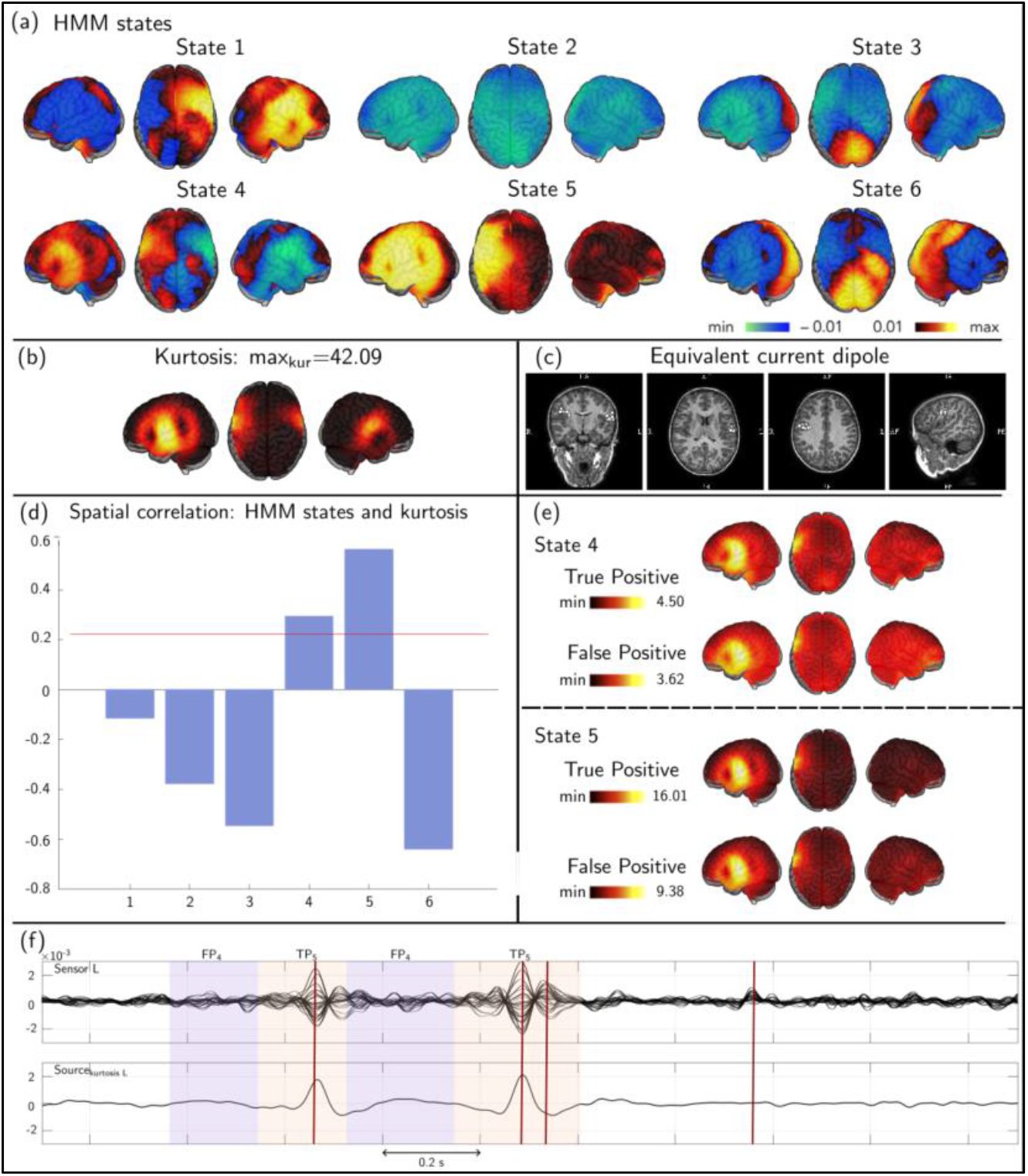
aHMM-based IED detection in patient 3. All is as in Fig. 5. Here, two epileptiform states were identified automatically by the kurtosis correlation test (states 4 and 5). Both states detect IED events at the left focus. Panel f: upper part, MEG data of left-hemisphere sensors; lower part, maximum-kurtosis source signal localized in the left centro-temporal part. Only VDS-IEDs coming from the left-hemisphere are seen. Note that IED detection events from two states (panel f; state 4, violet shade; state 5, orange shade) may not overlap in the aHMM pipeline.

**Figure 7.**
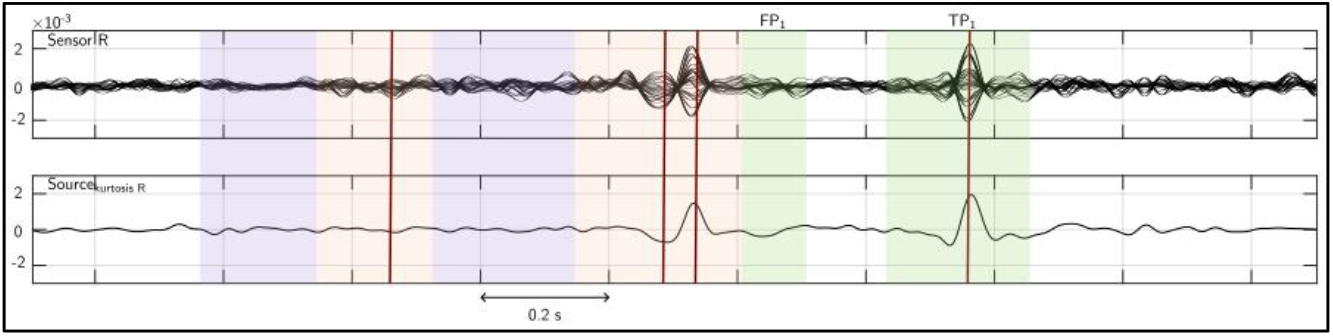
Visually selected aHMM state 1 of patient 3. Time course illustrated in the same 2-second period than in Fig.6(f), but this time coming from the right focus. Upper part, MEG data of right-hemisphere sensors; lower part, maximum-kurtosis value of in the right centro-temporal part. Signal shows IED detection events of state 1 (green shade) with the IED detection events of automatically selected states 4 (violet) and 5 (orange). Vertical lines indicate VDS-IEDs. VDS-IEDs coming from the left-hemisphere are not seen anymore (first and second VDS-IEDs in Figure 7). Contrary, the last VDS-IED coming from the right focus can now be seen clearly.

## 4. Discussion and conclusion

The development of a fully automatic method to detect and localize IEDs, both spatially and temporally, is an ambitious and long-standing goal in clinical epilepsy (Feys & De Tiège, 2023). While the visual analysis of MEG recordings in epileptic patients is certainly successful (Bagic et al., 2011; Laohathai et al., 2021), it remains a time-consuming process (on average ∼8h (De Tiège et al., 2017)) not devoid of subjective interpretation and limitations. Data-driven pipelines for reliable, fast, user-independent IED detection would represent an important step to move the field of clinical MEG forward by enabling subjectivity-free data analysis and harmonizing data analysis practices, which would be necessary for international multicenter studies (Bagić & Rampp, 2020). Here, we proposed and compared the efficiency of two distinct automatic pipelines—one based on ICA of MEG signals and the other based on aHMM of source-projected MEG signals—in a cohort of 10 school-age children with focal epilepsy. Both approaches proved similarly efficient and allowed fully unsupervised IED detection, at least in patients with unifocal epilepsy. In multifocal patients, the automated identification of epileptiform ICs/aHMM states tended to miss IED activity in some foci. However, bypassing this single step (i.e., spatial correlation test with kurtosis mapping) by a visual inspection of IC/aHMM state maps allowed to recover epileptiform activity from all epileptic foci. Importantly, both ICA- and aHMM-based pipelines successfully detected the large majority of IEDs identified visually by an experienced MEG clinician (again, at the price of a mild user-dependent input in multifocal epilepsy cases). What’s more, they allowed to reveal low-amplitude IEDs that were missed altogether during visual IED identification.

The building block of each pipeline consisted in a data-driven algorithm to extract distinct spatio-temporal patterns from MEG data. Our first pipeline relied on ICA, a blind source separation technique commonly used in MEG/EEG to identify physiological artefacts (Vigário, 1999) as well as functional connectivity networks (Brookes et al., 2011; Wens et al., 2014) and their metastable synchronization properties (Wens et al., 2019). Its ability to isolate epileptiform activity has been demonstrated repeatedly both in EEG (Hoeve et al., 2003; James et al., 1997) and MEG (Chirkov et al., 2022; Ossadtchi et al., 2004). The reason why ICA successfully isolates IED activity is the same than why kurtosis mapping successfully localizes IEDs: ICA is based on the optimization of a fourth-order cumulant statistic closely related to kurtosis (Hyvärinen et al., 2001), so IC signals tend have high kurtosis and to contain sporadic, high-amplitude events such as epileptic spikes (see also (Wens et al., 2019), for further discussion in the context of dynamic functional connectivity analysis). This leads to epileptiform ICs that correspond to high-kurtosis network activity and thus co-localize with the epileptic network imaged by kurtosis mapping. Compared to HMM, one limitation of ICA is that it does not provide a hard decision on the timing of IED events. This limitation may be circumvented in several ways, such as the use of a pre-defined detection threshold (Ossadtchi et al., 2004) or of clustering methods (Chirkov et al., 2022; Ossadtchi et al., 2005). Here, we used a 2-state HMM applied to the amplitude time course of each epileptiform IC in order to cluster signal time points and thus obtain an unsupervised detection of time windows containing IED events. That is important for practical purposes as the visual identification of spike timing events is indeed time consuming in routine clinical practice.

The building block of our second pipeline was a multi-state HMM applied to multivariate source amplitude MEG data. The HMM is a state clustering technique that finds many applications from speech processing (Rabiner et al., 1986; Rabiner, 1989) to biomedical applications (Penny et al., 1999; Flexer et al., 2002; Rezek et al., 2005), including the analysis of transient oscillatory power bursts and of the short-time activation dynamics of functional connectivity networks (Baker et al., 2014; Coquelet et al., 2022; Vidaurre et al., 2016, 2018). The ability of HMM to infer epileptic network states encompassing IEDs was tested recently with success (Seedat et al., 2022; Ye et al., 2022), at least using the TE-HMM implementation (Vidaurre et al., 2018). Here, we chose to build our HMM pipeline based on aHMM to automatically split MEG signals into periods of low and high amplitude. Our aHMM was able to identify one or more epileptiform states co-localizing with kurtosis mapping with an efficiency quite similar to ICA but also with similar issues regarding multifocal epilepsy and with a similar way out (i.e., selection of epileptiform states by a visual inspection of state maps). Of note, the HMM allowed to obtain a hard decision on the timing of IED event without the need of extra clustering steps (as with ICA). In our data, the HMM also turned out more efficient in one of the three patients with multifocal epilepsy. This being said, the HMM may suffer from test-retest reliability issues (Seedat et al., 2022). Given that both ICA and HMM optimization algorithms use random initial points, this question may actually be raised for both our pipelines. As a sanity check, we thus re-ran our IED detection pipelines multiple times on the same subjects in order to evaluate this aspect (data not shown). The ICA turned out to be very reliable and the outcome of our ICA-based IED detection pipeline was virtually identical each time. On the other hand, multivariate aHMM state classification appeared more volatile. In particular, a single epileptiform state would in some instances appear as two or more states (as observed by (Seedat et al., 2022) in the case of TE-HMM). This instability of the HMM inference is presumably due to the fact that we applied it at the individual level and with relatively short MEG recordings (5 min as all patients had frequent IEDs on previous scalp EEG), so it might be alleviated when using longer MEG recordings as recommended for MEG done in the context of the presurgical evaluation of refractory focal epilepsy (Bagic et al., 2011). Notwithstanding, the impact of this issue is eventually limited at the level of IED event detection and their statistics (IED-DR, sensitivity and specificity) because these events were constructed by gathering all epileptiform states (somewhat similarly to the idea of HMM metastates of (Vidaurre et al., 2017)).

Another crucial aspect of our study compared to previous works (Chirkov et al., 2022; Ossadtchi et al., 2004, 2005; Seedat et al., 2022; Ye et al., 2022) is that we compared ICA- and aHMM-based IED detection with the traditional process of visual inspection by an experienced MEG clinician blind to the outcome of both pipelines. Both approaches successfully recovered the large majority of VDS-IEDs, again at the price of abandoning full automatization in multifocal epilepsy. In the latter case though, the decision process of selecting which IC/aHMM state is epileptiform (based, e.g., on their spatial map) is arguably much easier and faster than the systematic process of manual IED identification throughout the whole MEG recording. Therefore, even though the two proposed approaches may not yet be fully automated, their usage may drastically simplify the workload for MEG clinicians and diminish interpretative ambiguities. Perhaps more importantly, our (semi)automated ICA and aHMM detection algorithms disclosed almost two times as much IED events than the clinician. We demonstrated that events missed out by visual identification co-localized with IEDs but exhibited much lower amplitudes of the order of background noise—which was the very reason why they were left undetected in the visual process. In the ICA pipeline, the ability to distinguish low-amplitude IEDs from noise was due to the denoising inherent to the isolation of IED activity into a small number of ICs. In the HMM pipeline, similar denoising may be due to the source projection step. That is, in our opinion, a crucial advantage of our IED detection methods.

One of the main limitations of automatic IED-identification pipelines in the context of the presurgical evaluation of refractory focal epilepsy is that they won’t be able to make an automatic distinction between true IEDs and physiological transients (e.g., Rolandic, supramarginal, occipital sharp transient activities) that may be confounded with IEDs as they share similar features (sharp slopes, duration, repetitive character, etc. (Laohathai et al., 2021; Rampp et al., 2020)). Such distinction will presumably require human expertise or, ultimately, entrained artificial intelligence, to classify the identified events as pathological or physiological based on their spatial, temporal or spectral dynamics. Also, the pipelines proposed in this study have been tested in a small population of children with focal epilepsy and frequent IEDs in previous scalp EEG. Most of the included patients had SeL-ECTS, which is characterized by IEDs with high signal to noise ratio and mainly perirolandic/perisylvian irritative zones. The high IED frequency and signal to noise ratio might partly explain the excellent sensitivity of the proposed approaches to automatically detect and localize IEDs. The next natural step is therefore to prospectively test those pipelines in consecutive patients with refractory focal epilepsy who will have various types of IEDs in terms of frequency (about 20-30% of such patients do not have IEDs (De Tiège et al., 2012; Duez et al., 2019; Knowlton et al., 2006; Rampp et al., 2019)) and locations. Ultimately, the comparison with post-operative outcome will also reveal their accuracy and predictive value of being seizure-free after surgical resection. Finally, the fact that the two proposed automatic approaches failed to identify the multiple irritative zones in some patients imposes that the analyses cannot be fully automated and requires some degree of human validation relying on visual reading of the MEG data.

In conclusion, we have developed, tested, and compared two IED detection pipelines, one based on ICA and the other based on aHMM of MEG data. Although they cannot be used in a fully automated way yet, they require a fairly mild user-dependent decision and decreases substantially the duration of the analysis process. Further, they appear to improve substantially the detection sensitivity to low-amplitude IEDs. As such, they represent a significant improvement over the currently established clinical practice.

## Author contributions

R.F.M., X.D.T. and V.W. designed study; O.F., E.J. and V.W. acquired data; R.F.M. and V.W. contributed to analytic tools; R.F.M., O.F., V.W. and X.D.T. analyzed data; R.F.M., O.F., E.J., A.A., C.U., V.W. and X.D.T. contributed to manuscript drafting or revision, X.D.T. obtained funding.

## Acknowledgments

R.F.M. is supported by a research grant of the Fonds Erasme (Brussels, Belgium; Research Convention: « Les Voies du Savoir II »). O.F. is supported by the Fonds pour la formation à la recherche dans l’industrie et l’agriculture (FRIA, Fonds de la Recherche Scientifique (FRS-FNRS), Brussels, Belgium).

X.D.T is Clinical Researcher at the FRS-FNRS (Brussels, Belgium).

The MEG project at the CUB – Hôpital Erasme is financially supported by the Fonds Erasme (Brussels, Belgium; Research Conventions “Les Voies du Savoir I & II”).

